# Cortical tension regulates Hippo signaling via Par-1-mediated Kibra degradation

**DOI:** 10.1101/2023.06.18.545491

**Authors:** Sherzod A. Tokamov, Stephan Buiter, Anne Ullyot, Gordana Scepanovic, Audrey Miller Williams, Rodrigo Fernandez-Gonzalez, Sally Horne-Badovinac, Richard G. Fehon

## Abstract

The Hippo pathway is an evolutionarily conserved regulator of tissue growth. Multiple Hippo signaling components are regulated via proteolytic degradation. However, how these degradation mechanisms are themselves modulated remains unexplored. Kibra is a key upstream pathway activator that promotes its own ubiquitin-mediated degradation upon assembling a Hippo signaling complex. Here, we demonstrate that Hippo complex-dependent Kibra degradation is modulated by cortical tension. Using classical genetic, osmotic, and pharmacological manipulations of myosin activity and cortical tension, we show that increasing cortical tension leads to Kibra degradation, whereas decreasing cortical tension increases Kibra abundance. Our study also implicates Par-1 in regulating Kib abundance downstream of cortical tension. We demonstrate that Par-1 promotes ubiquitin-mediated Kib degradation in a Hippo complex-dependent manner and is required for tension-induced Kib degradation. Collectively, our results reveal a previously unknown molecular mechanism by which cortical tension affects Hippo signaling and provide novel insights into the role of mechanical forces in growth control.

## Introduction

Elucidating the mechanisms that ensure robustness and reproducibility of organ size is a fundamental problem in cell and developmental biology. A key regulator of tissue growth in animals is the Hippo pathway. At its core, the Hippo pathway consists of a kinase cascade, composed of Ser/Thr kinases Hippo (Hpo) and Warts (Wts) and scaffold proteins Salvador (Sav) and Mats, that inhibits nuclear translocation of a pro-growth transcriptional co-activator, Yorkie (Yki). The activity of the kinase cascade is regulated by multiple upstream components, including a multivalent scaffold protein, Kibra (Kib). Kib localizes at the apical cell cortex of epithelial cells, where together with its binding partner, Merlin (Mer), it plays a key role in assembling and activating the Hippo kinase cascade to repress Yki activity. In the absence of a conventional receptor/ligand pair, however, little is known about how Kib activity is regulated.

A conserved feature of the Hippo pathway is its regulation via forces generated by F-actin and non-muscle myosin II (actomyosin). Actomyosin forces are regulated by the small GTPase RhoA (Rho1 in Drosophila), which can be activated or deactivated by guanine exchange factors (GEFs) or GTPase activating proteins (GAPs), respectively (Piekny et al., 2005). Rho1 potently stimulates actomyosin contractility by simultaneously activating Diaphanous, a formin that promotes linear F-actin bundle assembly, and Rho kinase (Rok), which phosphorylates and activates the myosin regulatory light chain (Lecuit et al., 2011). Studies over the years have shown that both F-actin polymerization and myosin activation stimulate the activity of Yki and its mammalian homologs, YAP/TAZ (Sansores-Garcia et al., 2011; Fernandez et al., 2011; Dupont et al., 2011; Wada et al., 2011; Aragona et al., 2013; Rauskolb et al., 2014; Ibar et al., 2018). Mechanistically, cytoskeletal tension was shown to promote Yki/YAP activity via LIM domain Ajuba family proteins, such as Ajuba (Jub), which accumulate at cell-cell junctions in response to tension and sequester Wts (Rauskolb et al., 2014; Pan et al., 2016; Ibar et al., 2018). However, this mechanism acts directly on the core kinase cascade, and it remains unclear whether or how tension could regulate the upstream Hippo signaling components, such as Kib.

A key determinant of signaling ouput downstream of Kib is its protein level. Loss of Kib results in tissue overgrowth via Yki activation, while the opposite occurs under ectopic Kib expression (Yu et al., 2010a; Genevet et al., 2010; Baumgartner et al., 2010; Su et al., 2017; Tokamov et al., 2021). Furthermore, upstream regulators, including Mer, Expanded and Kib, are transcriptionally upregulated by Yki, making their abundance a key component of a conserved negative feedback loop (Hamaratoglu et al., 2006; Genevet et al., 2010; Yee et al., 2019). Thus, control of Kib abundance provides an important point for modulating Kib-mediated Hippo pathway activation. In addition to transcriptional regulation, Kib abundance is regulated via ubiquitin-mediated proteolytic turnover (Tokamov et al., 2021). Upon assembly of the Hippo complex, Kib is ubiquitinated via the SCF^Slimb^ E3 ubiquitin ligase and subsequently degraded. This mechanism simultaneously requires a consensus recognition motif in Kib that recruits SCF^Slimb^, as well as the two N-terminal WW domains in Kib that mediate Hippo complex formation. Notably, Kib levels are elevated in cells with lower cortical tension, suggesting that actomyosin-generated forces could regulate Kib abundance (Tokamov et al., 2021). How Kib levels are modulated by mechanical forces and whether this mechanism involves additional mechanosensitive components remains unknown.

In this study, using genetic, osmotic, and pharmacological manipulations of myosin activity, we report that actomyosin-generated cortical tension modulates Kib abundance. We find that increasing myosin activity and cortical tension lowers Kib abundance, while acutely inhibiting myosin activity elevates Kib levels. Importantly, we find that tension-mediated Kib degradation occurs independently of the previously identified Jub-Wts mechanism. Instead, our results suggest that the serine/threonine kinase Par-1 promotes Kib degradation in a tension dependent manner. Together, our findings provide evidence that tension regulates upstream Hippo signaling and advance our understanding of how mechanical forces can influence the activity of a signaling pathway.

## Results

### Cortical tension promotes Kib degradation

Previously, we showed that Kib’s role in assembling the Hippo signaling complex promotes its own degradation, thereby forming a post-translational negative feedback loop (Tokamov et al., 2021). Additionally, our results suggested that this feedback mechanism is modulated by cortical tension because cell clones under compression display higher levels of Kib than their neighbors. To better understand how Kib abundance is regulated, we sought to genetically manipulate non-muscle myosin II (myosin) contractility as a means to affect cortical tension. To visualize Kib, we used Kib tagged with the green fluorescent protein (GFP) and expressed under the ubiquitin promoter (Ubi>Kib-GFP), a transgene that we previously showed to be an effective reporter of post-translational changes in Kib abundance (Tokamov et al., 2021). We first transiently ectopically expressed RhoGEF2 to activate Rho1 upstream of myosin in the posterior compartment of the wing imaginal disc using the *hh>Gal4* driver combined with ubiquitously expressed *Gal80^ts^* (*tub>Gal80^ts^*). Strikingly, expression of RhoGEF2 for 24h resulted in a significant decrease in Kib levels (Fig. 1A-B). Using phospho-specific antibody against the phosphorylated myosin regulatory light chain (pMRLC, Zhang and Ward, 2011), we confirmed that myosin activity was upregulated under these conditions (Fig. S1A-A’’). Additionally, consistent with the function of Kib in repressing Yki and the role of tension in promoting Yki transcriptional activity (Genevet et al., 2010; Yu et al., 2010b; Baumgartner et al., 2010; Rauskolb et al., 2014), we saw increased nuclear Yki accumulation upon RhoGEF2 expression (Fig. S1B-B’’). In contrast, the abundance of Ubi>Kib^ΔWW1^-GFP, a variant of Kib that is insensitive to Hippo complex-mediated degradation (Tokamov et al., 2021), was not affected by RhoGEF2 expression (Fig. 1C-D). These results are consistent with our previous observation that the abundance of wild-type Kib, but not Kib^ΔWW1^, was elevated in mosaic clones with decreased cortical tension (Tokamov et al., 2021) and suggest that cortical tension regulates Kib abundance via complex-mediated degradation.

**Figure 1:**
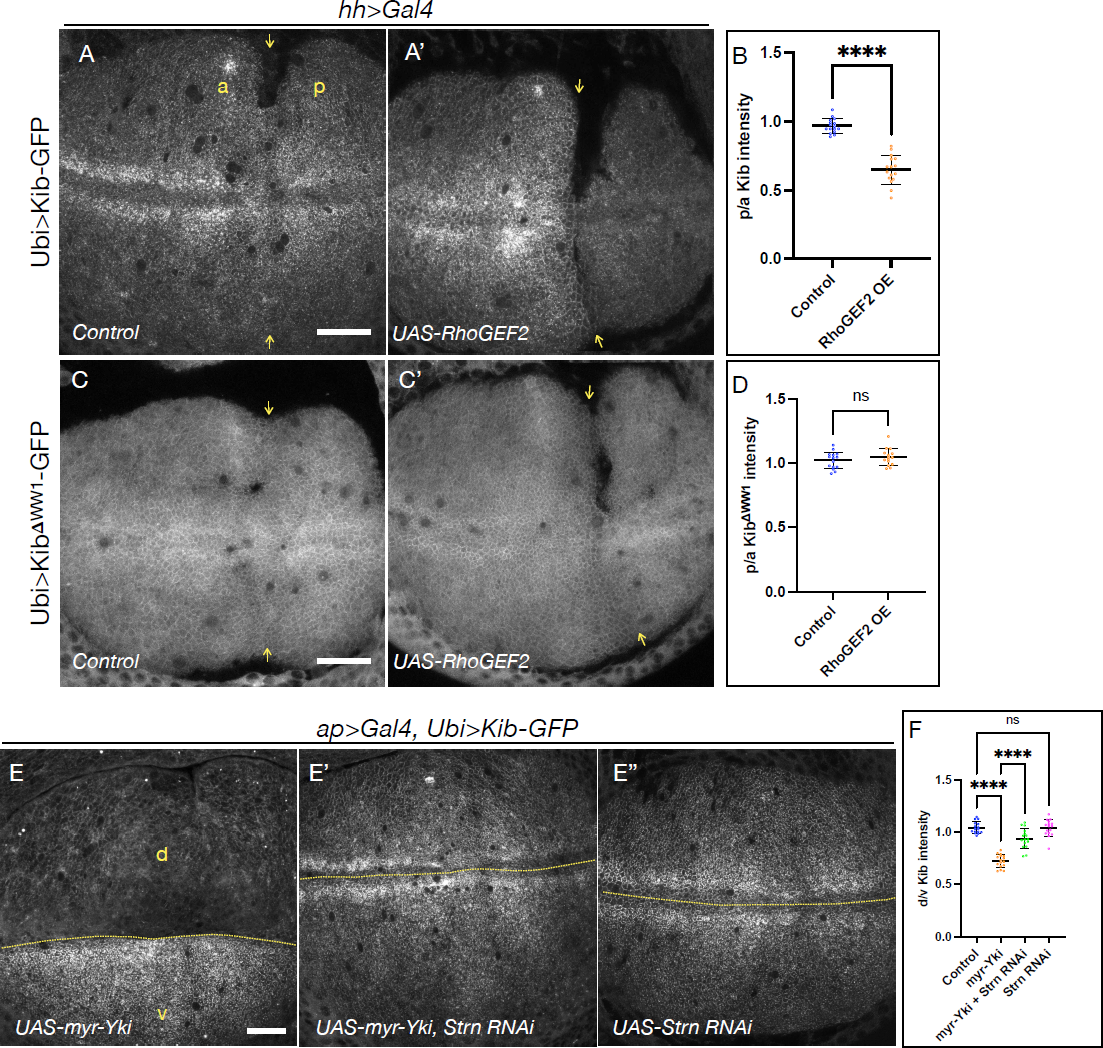
Cortical tension promotes Kib degradation. A-A’) Compared to control tissues (A), ectopic RhoGEF2 expression results in a significant decrease in Kib abundance (A’). Yellow arrows indicate anterior-posterior (a-p) boundary. B) Plot of p/a ratio of Ubi>Kib-GFP mean fluorescence intensity under the conditions shown in A-A’. C-C’) In contrast to wild-type Kib, Kib^!WW1^ is not affected by RhoGEF2 expression. D) Plot of p/a ratio of Ubi>Kib ^!WW1^-GFP mean fluorescence intensity under the conditions shown in C-C’. E-E’’) Ectopic myr-Yki expression leads to a significant decrease in Kib abundance (E), and depletion of Strn-MLCK suppresses the effect of myr-Yki expression (E’). Depletion of Strn-MLCK alone has no effect on Kib abundance (E’’). Dashed lines indicate the dorsal-ventral (d-v) boundary. Scale bars = 20μm. F) Plot of d/v ratio of Ubi>Kib-GFP mean fluorescence intensity under the conditions shown in E-E’’ (a representative control image of Ubi>Kib-GFP is shown in A). Unless indicated otherwise, statistical significance for two group comparisons was calculated using Mann-Whitney test. For more than two groups, One-way ANOVA followed by Tukey’s HSD test was used. Also, unless otherwise noted data are shown as the mean ± SD and significance values are represented as follows: ****p σ; 0.0001, ***p σ; 0.0001, **p σ; 0.001, *p σ; 0.05, ns = not significant.

We also previously showed that cortically localized Yki promotes myosin activation via a myosin light chain kinase called Stretchin (Strn-MLCK, Xu et al., 2018). Specifically, ectopic expression of myristoylated Yki (myr-Yki) leads to a significant increase in myosin activation. Therefore, we asked if myr-Yki expression also affected Kib levels. As with RhoGEF2 expression, ectopic myr-Yki expression substantially diminished Kib levels (Fig. 1E & F). Depletion of Strn-MLCK strongly suppressed the effect of myr-Yki, suggesting that this effect was mediated via myosin activity (Fig. 1E’, F). However, Strn-MLCK depletion alone did not increase Kib levels (Fig. 1E’’, F), so the physiological significance of this effect is unclear.

### Changes in osmotic conditions alter myosin activity and cortical tension

Our genetic manipulations of cortical tension support the model that tension promotes Kib turnover. We sought to further test this model by acutely manipulating myosin activity and cortical tension. Changes in osmotic pressure have been used extensively to alter cortical cytoskeletal organization and tension (Di Ciano et al., 2002; Guilak et al., 2002; Sinha et al., 2011; Boulant et al., 2011; Stewart et al., 2011; Pietuch et al., 2013; Diz-Muñoz et al., 2016; Roffay et al., 2021). Therefore, we decided to subject wing imaginal discs to different osmotic conditions (see Methods) and examine the effect on myosin activity and cortical tension. We aimed to alter osmolarity gently enough to avoid cell death or tissue rupture and ensure reversibility of the effected changes.

We first examined the effect of different osmotic conditions on myosin activity using the anti-pMRLC antibody. Compared to isotonic controls, we saw a slight decrease in pMRLC staining under hypertonic conditions, although this effect was not statistically significant, likely due to the background associated with the antibody (Fig. 2A-B’, D). Under hypotonic conditions, however, we observed a substantial increase in pMRLC (Fig. 2C-D), indicating that osmotic manipulations can acutely alter myosin activation. To ask if the observed changes in myosin activity affected cortical tension, we performed laser-cutting experiments on individual bicellular junctions and measured their initial recoil velocity. Because of the inherent heterogeneity of cell junctional lengths and apical areas across the wing imaginal epithelium, we restricted all laser cuts to the ventral-anterior region of the presumptive wing blade, which normally contains cells with larger apical areas and longer junctions. Additionally, we were careful to avoid proximity to mitotic cells (Fig. S2A). Consistent with the observed differences in pMRLC staining, we saw a significant decrease in recoil velocities under hypertonic conditions and an even more significant increase in recoil upon hypotonic shift (Fig. 2E-F, Movie 1). We observed no correlation between junctional length and recoil velocities (Fig. S2B). Collectively, these results show that osmotic manipulations can be used to acutely modulate myosin activity and cortical tension in the imaginal epithelium.

**Figure 2:**
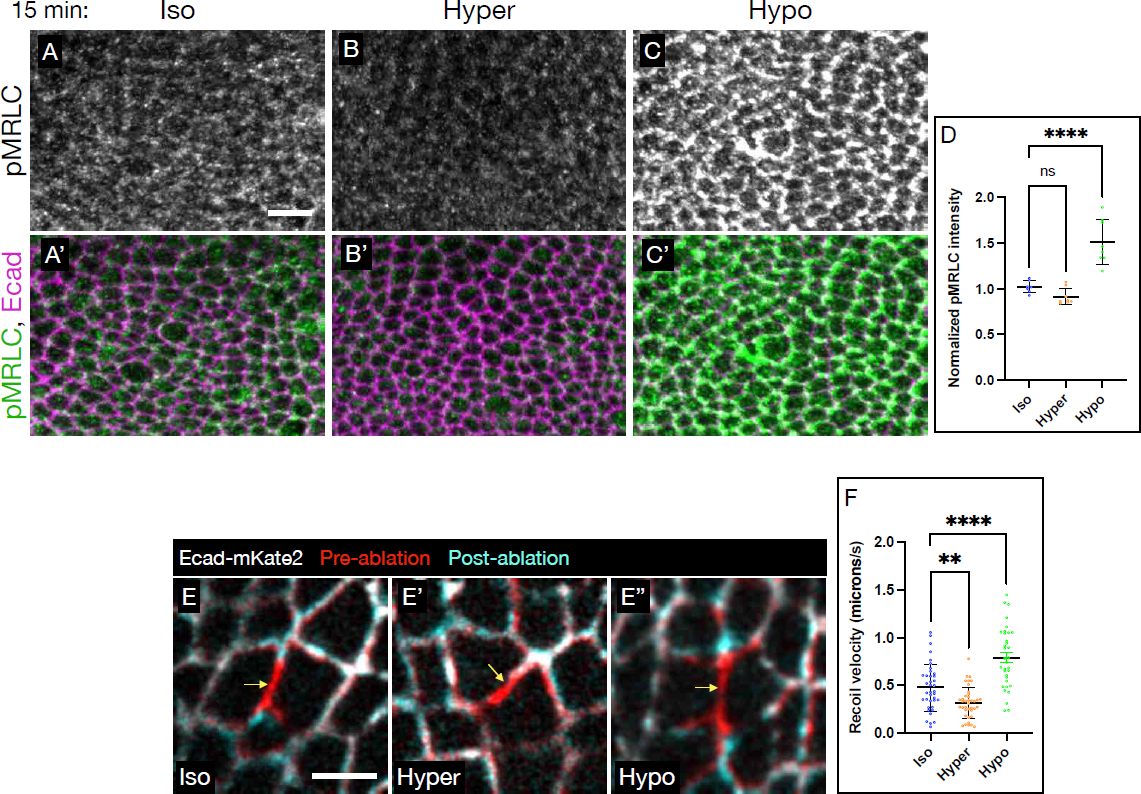
Changes in osmotic conditions alter myosin activity and cortical tension. A-C) Compared to isotonic conditions (A-A’), hypertonic conditions lead to a slight decrease in pMRLC (B-B’), whereas hypotonic conditions significantly increase pMRLC signal (C-C’). Scale bar = 5μm. D) Quantification of mean pMRLC intensity under osmotic conditions shown in A-C. E-E’’) Representative overlay images of Ecad-mKate2 in wing imaginal disc cells pre-(red) and post-ablation of bicellular junctions (arrows) under isotonic (E), hypertonic (E’), hypotonic E’’) conditions. Scale bar = 3μm. F) Quantification of initial junction recoil velocities measured from the ablation experiments.

To further validate that osmotically-altered cortical tension can induce biologically relevant consequences, we once again examined the localization of Yki-YFP. Because tension is known to promote nuclear Yki accumulation, we hypothesized that hypertonic conditions (lower tension) should lead to less nuclear Yki, while hypotonic conditions should result in more nuclear Yki. For these experiments, in order to see a definitive effect on Yki localization, we increased the incubation time to 30 min. Consistent with our hypothesis, shifting tissues from an isotonic to a hypertonic solution resulted in a decrease in nuclear Yki (Fig. S3A-B’’, D). Conversely, shifting to a hypotonic solution led to a dramatic increase in nuclear Yki (Fig. S3C-D). To further demonstrate the significance of this effect, we decided to first concentrate Yki in the nuclei under hypotonic conditions and then shift to a hypertonic environment. Hypotonically-induced nuclear Yki accumulation was completely reversed after incubation in a hypertonic solution (Fig. S3E-H). Together, these results show that osmotic manipulations can be used to acutely modulate myosin activity and cortical tension in an epithelial monolayer with biologically relevant consequences.

### Osmotic shifts affect Kib abundance

Given our result that changes in the medium osmolarity can influence cortical tension, we asked if the osmotic manipulations also affected Kib abundance. Compared to isotonic conditions, incubating tissues in a hypertonic solution resulted in an increased Kib abundance at the apical cortex (Fig. 3A, B, & D). Moreover, Kib fluorescence also increased in more basal tissue sections, suggesting that the apical increase in Kib was not due to apical recruitment from the cytoplasmic pool (Fig. 3E). Conversely, in tissues incubated in a hypotonic solution, Kib abundance decreased dramatically both at the apical cortex and in more basal tissue sections compared to isotonic controls (Fig. 3C-E). In contrast, fluorescence intensity of Ubi>Kib^ιΔ1WW1^-GFP did not change significantly under osmotic shifts (Fig.S4A-C), suggesting that osmotically-induced Kib degradation is mediated via the previously described Hippo complex-dependent mechanism (Tokamov et al., 2021). These results demonstrate that cortical tension can rapidly modulate Kib abundance.

**Figure 3:**
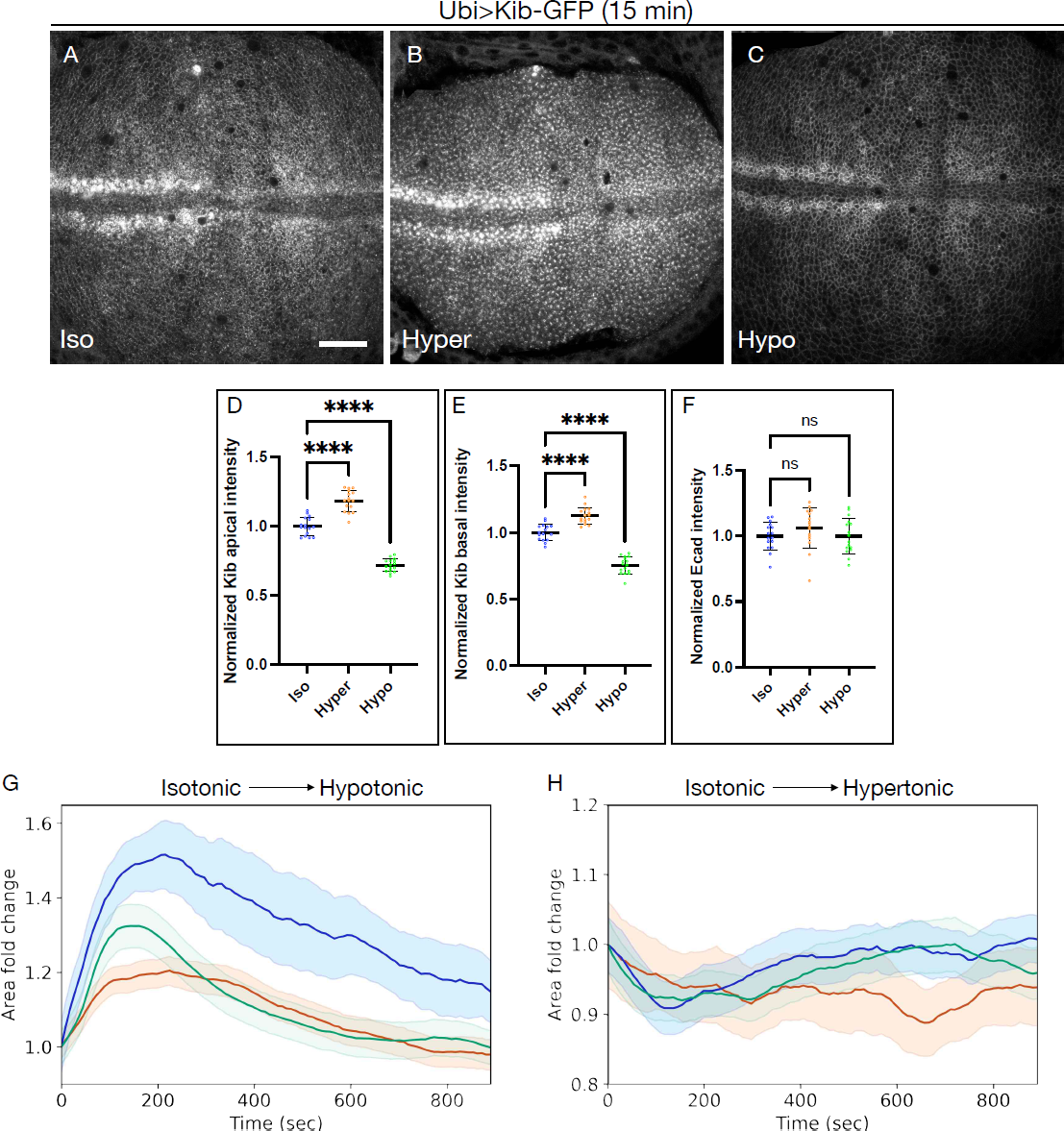
Osmotic shifts affect Kib abundance. A-C) Compared to isotonic conditions (A), hypertonic shift results in higher Kib abundance (B), while hypotonic shift leads to decreased Kib abundance (C). Scale bar = 20μm. D-F’) Quantification of apical Kib-GFP (D), basal Kib-GFP (E), and apical Ecad-mKate2 (F) mean fluorescence. G & H) Plots of cell area changes (mean ± SEM) obtained from tissues shifted from isotonic to hypotonic (G) or isotonic to hypertonic (H) conditions. Each curve represents an independent trial/tissue, with three tissues imaged for each condition.

We also considered the possibility that changes in apical area could alter Kib distribution and fluorescence intensity by concentrating or diluting the protein at the cortex. To address this point, we first measured fluorescent intensity of a junctional marker, Ecad-mKate2, and saw no significant changes under osmotic shifts (Fig. 3F). Next, we measured apical cell areas upon shifting wing imaginal tissues from isotonic to hypotonic or hypertonic conditions. Immediately after the addition of the hypotonic solution, apical areas increased rapidly during the first ∼3 min of incubation (Fig. 3G). However, this initial increase was followed by a steady decrease of areas in the next 12 min, until on average the areas returned roughly to their initial values (Fig. 3G, Movie 2). Similarly, addition of the hypertonic solution resulted in a transient reduction in apical areas during the first ∼3 min, followed by a steady return to their starting values by 15 min of incubation (Fig. 3H, Movie 3). Because we measured Kib fluorescence after 15 min of incubation (Fig. 3A-C), changes in apical area are unlikely to explain the observed changes in cortical Kib abundance under osmotic shifts. Collectively, these results suggest that changes in myosin activity may be responsible for changes in Kib abundance under osmotic shifts.

## Inhibition of myosin activity blocks Kib degradation under hypotonic conditions

If myosin-generated cortical tension is responsible for the observed decrease in Kib abundance under hypotonic shift, then inhibiting myosin activity under these conditions should block Kib degradation. To test this idea, we used Y-27632, a pharmacological inhibitor of Rho kinase (Rok) activity (Uehata et al., 1997). Under isotonic conditions, we observed a slight reduction in pMRLC staining when tissues were treated with Y-27632 (Fig. 4A-B’), though this effect was not statistically significant (Fig. 4I; again, possibly obscured by high antibody background staining). However, Y-27632 potently inhibited myosin hyperactivation induced by hypotonic conditions (Fig. 4C-D’, I).

**Figure 4:**
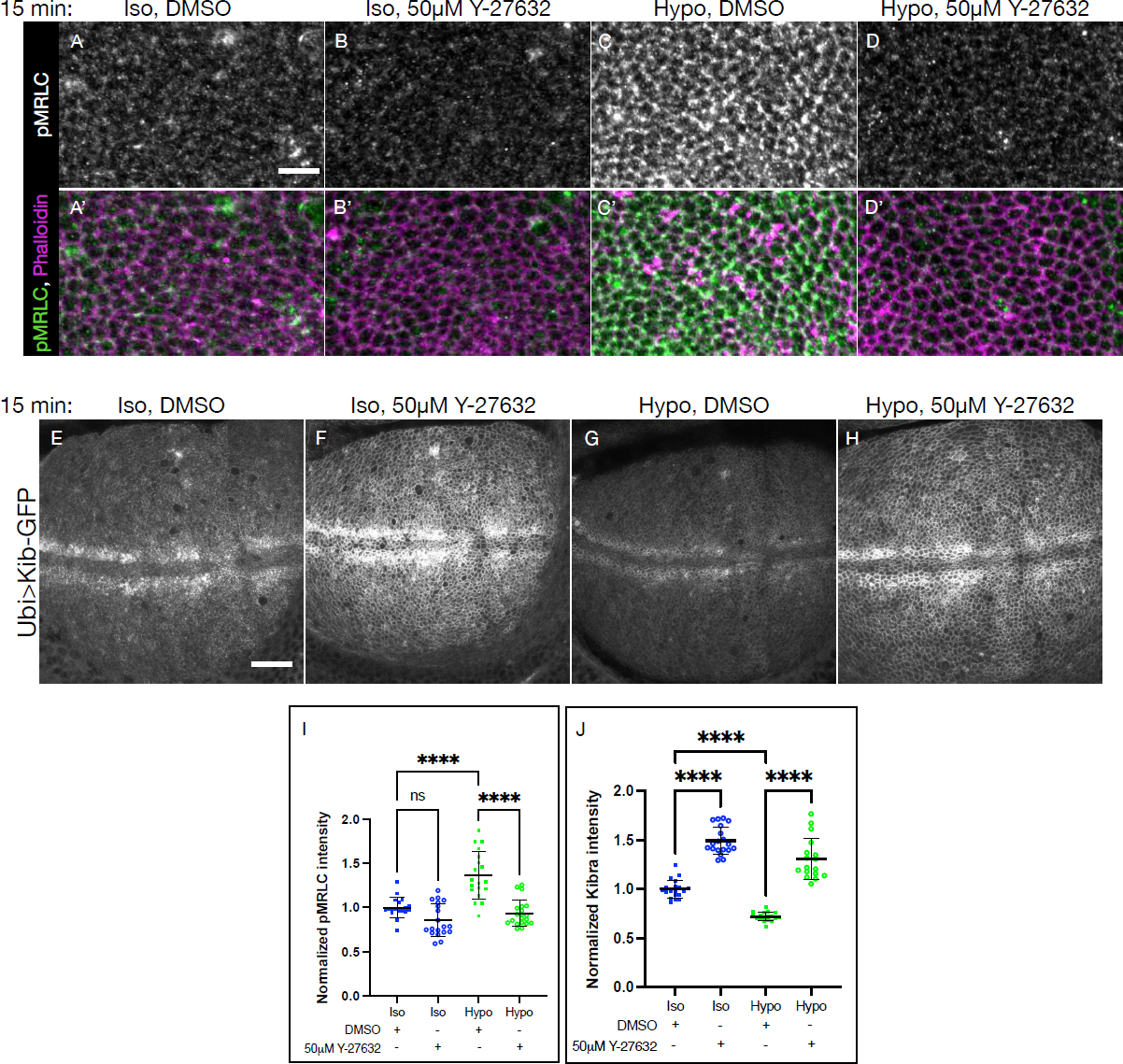
Treatment with Y-27632 raises Kib abundance and reverses the effect of hypotonic shift. A-D’) Staining with the antibody against pMRLC shows that Y-27632 treatment leads to a slight decrease in pMRLC staining under isotonic conditions (A-B’) and a blocks pMRLC upregulation under hypotonic shift (C-D’). Scale bar = 5μm. E-H) Compared to control (E), treatment with Y-27632 dramatically increases Kib abundance under isotonic conditions (F). Addition of Y-27632 also blocks the decrease in Kib abundance induced by hypotonic shift (G & H). Scale bar = 20μm. I) Plot of normalized mean pMRLC intensities obtained from experiments represented in A-D’. J) Plot of normalized mean Kib-GFP intensities obtained from experiments represented in E-H.

We next asked if Y-27632 treatment would affect Kib abundance. Strikingly, we observed a substantial increase in Kib abundance after 15 min of Y-27632 treatment under isotonic conditions (Fig. 4E, F, & J), supporting the idea that myosin activity promotes Kib degradation. To test if Rok inhibition could block the effect of hypotonic conditions, we incubated tissues in a hypotonic solution containing Y-27632. Addition of Y-27632 strongly blocked the effect of the hypotonic medium on Kib abundance (Fig. 4G, H, & J), suggesting that the effect of osmotic shifts on Kib abundance is caused by myosin activity.

We also considered the possibility that the effect of Y-27632 could be mediated via inhibition of another known target of this inhibitor, the atypical protein kinase C (aPKC). To test this possibility, we took advantage of a conditional allele of aPKC, *aPKC^as4^* that can be acutely inhibited using 1NA-PP1, an allele-specific analog of a potent kinase inhibitor (Hannaford et al., 2019a). As expected, under normal conditions, aPKC^as4^ was enriched at the apical cortex in the wing imaginal epithelium (Fig. S5A-A’). In contrast, treatment with 1NA-PP1 severely inhibited aPKC cortical localization (Fig. S5B-B’). On the other hand, treatment with Y-27632 did not significantly affect aPKC cortical localization in wild type imaginal discs (Fig. S5C-D’). Additionally, we treated wing discs homozygous for *aPKC^as4^* allele with 1NA-PP1 and did not observe any changes in Kib abundance upon aPKC inhibition (Fig. S5E-G). These results show that the effect of Y-27632 on Kib abundance is not due to aPKC inhibition and instead suggest that Kib degradation under hypotonic conditions is mediated via cortical tension.

### Par-1 regulates Kib abundance

How can tension influence abundance of a signaling protein such as Kib? Previous work identified a mechanism whereby tension inhibits Hippo signaling via Jub-mediated sequestration and inactivation of Wts/LATS at the cell-cell junctions (Rauskolb et al., 2014; Ibar et al., 2018). Surprisingly, although we observed a significant increase in myosin activity under hypotonic conditions (Fig. 2C), we did not detect significant changes in cortical Jub (Fig. S6A-D). Additionally, loss of Jub does not affect Kib levels (Fig. S6E). These observations suggest that cortical tension regulates Kib abundance independently of Jub.

Recently, cortical tension was shown to promote cortical localization of Par-1 in the Drosophila oocyte (Doerflinger et al., 2021). Interestingly, Par-1 mediates proteolytic degradation of an SCF^Slimb/βTrCP^ substrate Oskar (Morais-de-Sá et al., 2013) and is known to physically associate with Hippo signaling components and regulate tissue growth (Huang et al., 2013, p. 1). We therefore hypothesized that cortical tension could promote Kib degradation via Par-1. To test this idea, we first asked if Par-1 regulates Kib abundance using Ubi>Kib-GFP as a reporter. Kib levels increased significantly upon Par-1 depletion (Figs. 5A & C, S7A), whereas ectopic Par-1 expression resulted in a significant reduction in Kib levels (Fig. 5B & C). In contrast, loss of Par-1 did not affect the abundance of Kib^ΔWW1^ (Fig. 5D & E), suggesting that Par-1 regulates Kib levels via Kib-mediated Hippo complex assembly (Tokamov et al., 2021).

**Figure 5.**
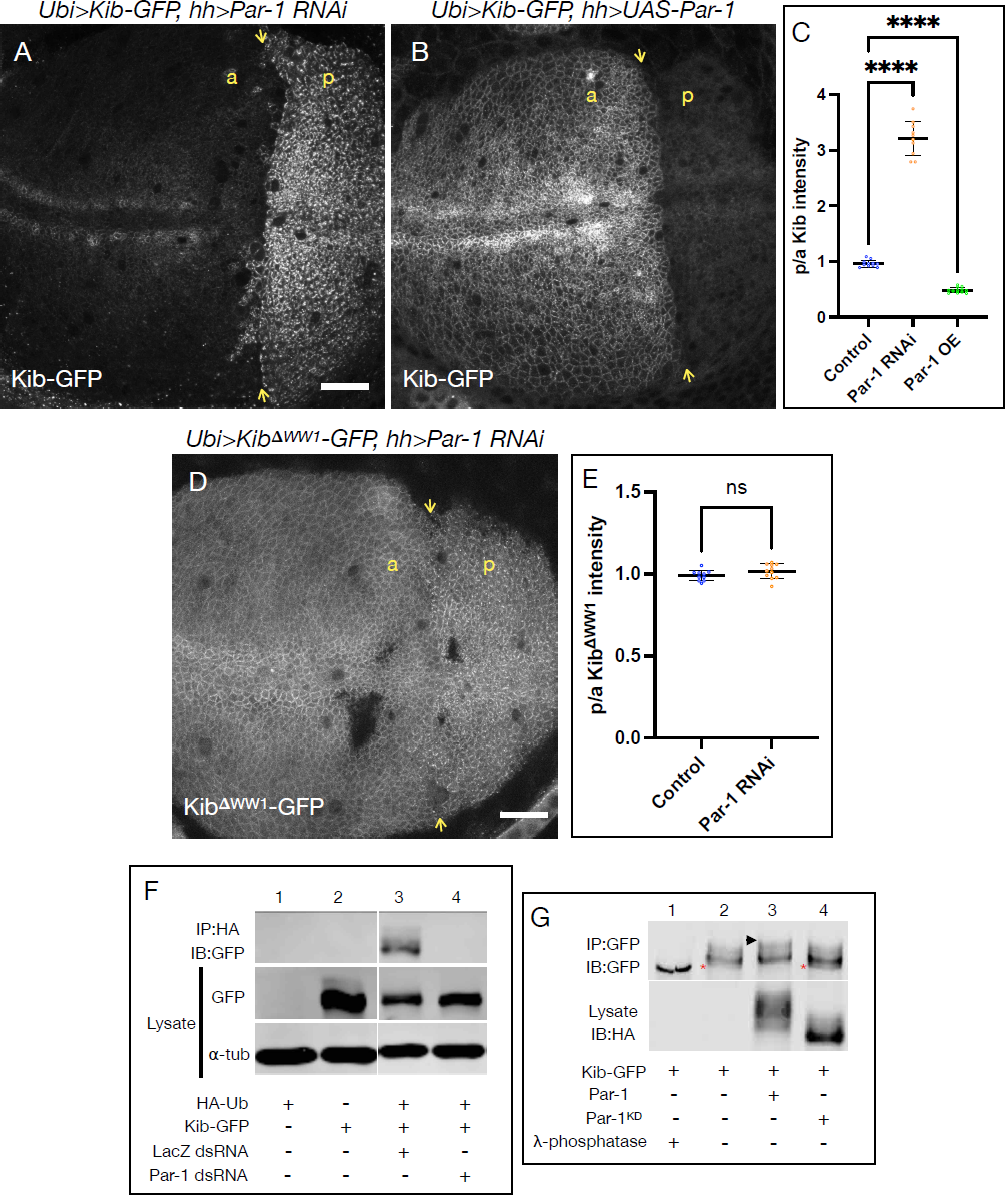
Par-1 promotes ubiquitin-mediated Kib degradation. A) Loss of Par-1 in the posterior wing disc compartment leads to a dramatic increase in Kib abundance. Yellow arrows indicate the anterior-posterior (a-p) boundary. Scale bar = 20μm. B) Ectopic Par-1 expression in the posterior wing disc compartment results in a strong decrease Kib abundance. C) Quantification of loss or gain of Par-1 function on Kib abundance as a posterior/anterior (p/a) ratio of mean fluorescence intensity. D) Loss of Par-1 in the posterior wing disc compartment does not affect Kib^ι1WW1^ abundance. E) Quantification of the effect of Par-1 depletion on Kib^ι1WW1^ abundance as a p/a ratio of mean fluorescence intensity. F) Kib immunoprecipitated from cultured S2 cells is normally ubiquitinated when cells are treated with a control LacZ dsRNA (lane 3). Depletion of Par-1 with dsRNA results in decreased Kib ubiquitination (lane 4). G) Treatment of Kib samples immunoprecipitated from S2 cells with λ-phosphatase results in non-phosphorylated, faster migrating band (lane 1) relative to phosphorylated, slower migrating bands observed without λ-phosphatase treatment (lane 2). Expression of wild-type Par-1, but not kinase-dead Par-1, leads to increased Kib phosphorylation (lanes 3 & 4). Asterisks indicate non-phosphorylated Kib pool, which disappears upon Par-1 expression. Arrowhead indicates a slower migrating (more phosphorylated) Kib pool.

Par-1 was previously shown to promote Hpo phosphorylation on Ser30, which inhibits Hpo activity (Huang et al., 2013). Therefore, we considered the possibility that the increase in Kib abundance under loss of Par-1 was indirectly caused by the lack of Hpo phosphorylation on Ser30. If this were the case, we reasoned that ectopic expression of Hpo construct that is refractory to Par-1 phosphorylation (Hpo^S30A^), would also lead to increased Kib abundance. To this end, we first transiently expressed wild-type Hpo in the posterior compartment of the wing imaginal disc and examined the effect on Kib levels. Consistent with our previous finding that Hpo promotes Kib ubiquitination and degradation (Tokamov et al., 2021), Kib levels decreased significantly under ectopic Hpo expression (Fig. S7B). Importantly, ectopic expression of Hpo^S30A^ also induced a decrease in Kib abundance (Fig. S7C), suggesting that the effect of Par-1 on Kib abundance is not mediated through Hpo phosphorylation.

We have previously shown that Kib is ubiquitinated via SCF^Slimb/βTrCP^ in cultured Schneider’s 2 (S2) cells. Because Par-1 is a kinase and is known to function with SCF^Slimb/βTrCP^, we asked if Par-1 affects Kib ubiquitination or phosphorylation in S2 cells. We observed that while Kib was ubiquitinated in control cells, Par-1 depletion led to a strong decrease in Kib ubiquitination (Fig. 5F). To test if Par-1 promotes Kib phosphorylation, we performed gel-shift assays in the presence of wild type or kinase-dead Par-1 (Par-1^KD^). Kib is normally phosphorylated in S2 cells, as it appears as a smeary, slow migrating band that collapses into a single, faster migrating band with phosphatase treatment (Fig. 5G, Tokamov et al., 2021). Addition of active Par-1, but not Par-1^KD^, resulted in an upward shift of Kib band and disappearance of the lower, non-phosphorylated pool, suggesting that Par-1 promotes Kib phosphorylation (Fig. 5G). Together, these results suggest that Par-1 regulates ubiquitin-mediated Kib degradation, possibly in a phosphorylation-dependent manner, and that tension could modulate Kib abundance via Par-1.

### Cortical tension promotes Par-1 association with the cortex

Given the previous report that myosin activity promotes cortical Par-1 localization in the *Drosophila* oocyte (Doerflinger et al., 2021), we wondered if a similar effect could be observed in the wing imaginal epithelium. To this end, we first examined Par-1 localization in control cells or in cells overexpressing RhoGEF2. Although known as a basolateral component, in the Drosophila blastoderm Par-1 also localizes at the apicolateral cortex (Bayraktar et al., 2006). In the wing imaginal disc, Par-1 displayed apical and basolateral localization, though its localization appeared diffuse in both places (Fig. 6A & A’). Transient overexpression of RhoGEF2 resulted in sharper Par-1 localization at the cell cortex both apically and basolaterally (Fig. 6B & B’), suggesting that tension can affect Par-1 cortical association. As an alternative approach, we also examined Par-1 localization using Airyscan confocal microscopy under hypotonic conditions. Similar to ectopic RhoGEF2 expression, Par-1 was more tightly associated with the apical and basolateral cortex under hypotonic compared to isotonic conditions (Fig. 6C-D’). These results suggest that tension promotes Par-1 association with the cell cortex.

**Figure 6:**
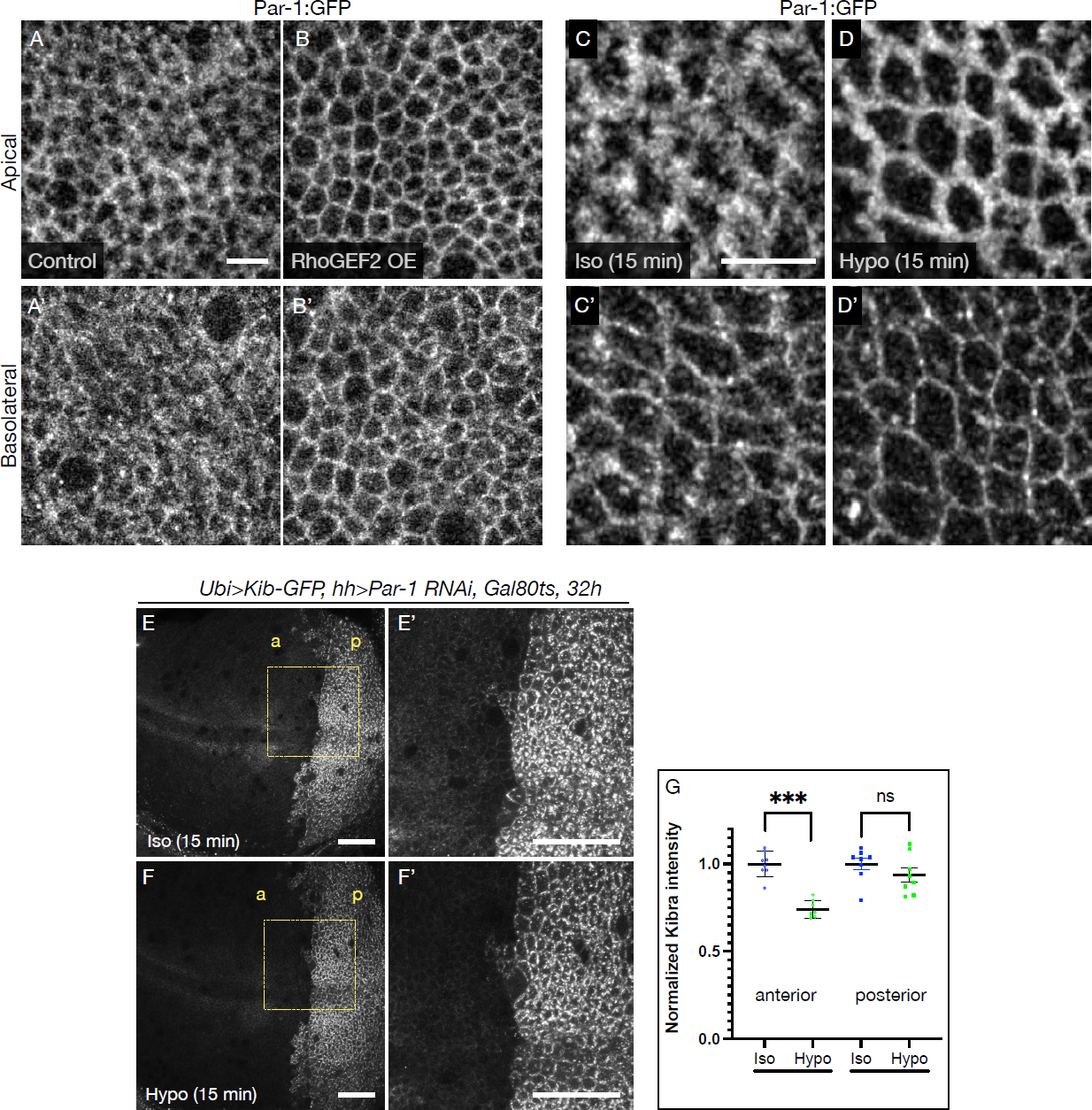
Cortical tension regulates Kib abundance by modulating cortical Par-1 association. A & B’) Compared to control cells (A-A’), ectopic RhoGEF2 expression leads to tighter Par-1 association with the apical and basolateral cell cortex (B-B’). Scale bar = 5μm. C & D’) Compared to isotonic conditions (C-C’), Par-1 becomes more tightly associated with the apical and basolateral cell cortex upon hypotonic shift (D-D’). Scale bar = 5μm. E-G) Under isotonic conditions, depletion of Par-1 in the posterior compartment (p) leads to a significant increase in Kib levels (E & E’, G). Shift to hypotonic conditions leads to a significant decrease in Kib abundance in the anterior region, but only a mild (not statistically significant) decrease in the posterior region where Par-1 was depleted (F & F’, G). Scale bars = 20μm.

### Par-1 is required for tension-dependent degradation of Kib

We reasoned that if tension regulates in Kib abundance via Par-1, then in the absence of Par-1 tension would have no effect on Kib levels. To test this idea, we depleted Par-1 in the posterior compartment of the wing imaginal disc using the *hh>Gal4* driver and quantified changes in Kib levels when tissues were shifted from isotonic to hypotonic conditions. While we observed a significant decrease in Kib intensity in the anterior (control) compartment, there was no significant change in the posterior (Par-1 depleted) compartment (Fig. 6E-G). Collectively, these results are consistent with the idea that tension modulates Kib abundance by controlling Par-1 cortical association.

## Discussion

Elucidating the molecular mechanisms that control the abundance of individual signaling components has been central to our understanding how the Hippo pathway is regulated (Ribeiro et al., 2014; Bosch et al., 2014; Rodrigues-Campos and Thompson, 2014; Aerne et al., 2015; Ma et al., 2018; Fulford et al., 2019; Wang et al., 2019; Misra and Irvine, 2019; Tokamov et al., 2021). However, how these different mechanisms themselves are regulated has been a poorly understood and intriguing area of investigation. Our work reveals a previously unrecognized role of cortical tension in modulating the abundance of a key upstream Hippo pathway component, Kib. Using a combination of genetic, osmotic, and pharmacological approaches, we demonstrate that cortical tension promotes Kib degradation. Our study also implicates Par-1 in mediating Kib degradation downstream of cortical tension. This work advances our understanding of how tension regulates Hippo signaling and provides novel insights into the role of mechanical forces in tissue growth and patterning.

We provide several lines of evidence that Kib is degraded as a result of myosin-generated tension. First, we show that activation of myosin via two different myosin light chain kinases, Rok (in the case of ectopic RhoGEF2 expression and Y-27632 treatment) and Strn-MLCK (in the case of ectopic myr-Yki expression) decreases Kib abundance (Figs. 1 & 4), suggesting that Kib degradation occurs irrespective of the upstream mechanisms of myosin activation. Second, although RhoGEF2 is known to promote both myosin activity and formin-mediated F-actin assembly, we did not observe a detectible effect on F-actin under transient RhoGEF2 expression, whereas myosin activity was substantially upregulated (Fig. S1). Third, we develop a simple, non-genetic and acute approach to dramatically increase myosin activity using hypotonic conditions and demonstrate that myosin activation using this method results in Kib degradation (Figs. 2-4). Thus, we propose that Kib degradation is modulated by cortical tension downstream of myosin activity.

Our results also suggest that Kib degradation downstream of cortical tension is mediated via Hippo complex formation. We have shown previously that Kib targets itself for ubiquitin-mediated degradation upon Hippo complex assembly, and this process requires Kib’s WW1 domain and a consensus degron motif recognized by SCF^Slimb/βTrCP^ (Tokamov et al., 2021). In this study, we observe that the abundance of wild-type Kib, but not Kib^ΔWW1^, is modulated via myosin activity, raising the question of how tension can promote complex-dependent Kib turnover. In the simplest scenario, cortical tension could promote the association of existing Kib complexes with a tension-sensitive component capable of modulating Kib levels. We identify Par-1 as a potential link in this process. Cortical tension was previously reported to promote cortical Par-1 localization in the *Drosophila* oocyte (Doerflinger et al., 2021). Here, we show that Par-1 promotes turnover of wild-type Kib, but not Kib^ΔWW1^, in vivo and regulates Kib phosphorylation and ubiquitination in cultured cells (Fig. 5). Importantly, cortical tension fails to induce Kib turnover in cells depleted of Par-1 (Fig. 6). Based on these results, and our observations that Par-1 becomes more tightly associated with the cell cortex under increased cortical tension (Fig. 6), we propose that cortical tension could regulate Kib levels at least in part by modulating the association of Par-1 with Kib signaling complexes. Interestingly, a recent study in human cells also suggested that tension generated at the ECM promotes degradation of a Hpo homolog, MST2, by modulating the physical interaction between MST2 and SCF^Slimb/βTrCP^ complex (Fiore et al., 2022). Thus, in a more complex scenario, cortical tension could also regulate Kib association with multiple components of the degradation machinery, including _SCFSlimb/βTrCP._

This study also highlights the recurring role of Par-1 in the regulation of Hippo signaling. Previous work suggested that Par-1 phosphorylates Hpo, which prevents Hpo association with Sav (Huang et al., 2013). Here, we find that Par-1 regulates Kib abundance independently of Hpo phosphorylation. Our observation that Par-1 regulates the abundance of wild-type Kib but not Kib^ιΔ1WW1^ suggests that Par-1 regulates Kib via the Hippo complex-mediated degradation mechanism involving SCF^Slimb/βTrCP^ (Tokamov et al., 2021). This idea is supported by a previous report that Par-1 functions with SCF^Slimb/βTrCP^ in promoting turnover of Oscar during Drosophila oocyte development (Morais-de-Sá et al., 2013). Notably, active Par-1 was also found to be a target of SCF^Slimb/βTrCP^-mediated degradation (Lee et al., 2012), further highlighting the tight association between Par-1, SCF^Slimb/βTrCP^, and Kib. It is still unclear what happens to the rest of the Hippo signaling complex after Kib is degraded, but the multifaceted function of Par-1 in inhibiting Hpo activity, Hpo-Sav interaction, and promoting Kib degradation could provide the means of dissociating the entire signaling complex upon Kib turnover.

One of the surprising observations in our study is that while cortical tension induced by hypotonic conditions triggers Kib degradation, it does not affect junctional Jub accumulation. Previous work has shown that increasing myosin activity via genetic manipulations enhances Jub recruitment to the adherens junctions, where Jub sequesters Wts (Rauskolb et al., 2014). This recruitment is thought to be mediated via tension-induced opening of α-catenin conformation, which exposes a region in α-catenin that binds Jub (Rauskolb et al., 2014; Alégot et al., 2019; Sarpal et al., 2019). Combined with these reports, our observations suggest that hypotonically-induced myosin activation may differ from that achieved via common genetic manipulations. Most obviously, this difference could be temporal – hypotonic treatment of wing imaginal discs induces dramatic myosin activation and Kib degradation within 15 minutes (Fig. 2), whereas genetic manipulations are done in the order of hours or days. Thus, α-catenin-mediated recruitment of Jub could occur at a longer timescale compared to the more dynamic Kib turnover. Counter to this argument, a recent study observed rapid junctional accumulation of Jub-Wts clusters at sites of high tension in the Drosophila developing notum (López-Gay et al., 2020). An alternative explanation is that hypotonically-induced myosin activation does not result in α-catenin conformational changes. The opening of α-catenin conformation at bicellular junctions is thought to be triggered via actomyosin forces applied orthogonally to the junctions (Rauskolb et al., 2019; López-Gay et al., 2020). This is unlikely to be the case under hypotonic conditions, where actomyosin generated forces would be generated parallel to the lateral membrane to counteract the mechanical stretching induced by hydrostatic pressure.

Finally, our study raises the question of the biological importance of tension-dependent Kib degradation. In the growing wing imaginal epithelium, cortical tension was shown to be higher at the tissue periphery than at the tissue center (LeGoff et al., 2013; Mao et al., 2013). Greater tension at the periphery was proposed to drive growth, possibly via Yki activity, to compensate for the lower concentration of growth-stimulating morphogens, which diffuse from narrow stripes of cells at the center of the tissue (Aegerter-Wilmsen et al., 2007, 2012; Hariharan, 2015; Pan et al., 2016). Additionally, we have previously reported that Kib degradation occurs more prominently at the tissue periphery, where cortical tension is higher (Tokamov et al., 2021). Based on these studies, we propose that tension-mediated Kib degradation could serve to pattern Hippo signaling activity, and therefore growth, across a growing epithelium such as the wing imaginal disc. How Kib degradation and other tension-regulated mechanisms, such as the Jub-Wts mechanism, are coordinated to control the Hippo pathway and growth remains unanswered. However, we believe this study provides valuable insights for future investigation of the molecular mechanisms by which mechanical forces affect tissue growth.

## Methods

### Drosophila husbandry

*Drosophila melanogaster* was cultured using standard techniques at 25°C (unless otherwise noted). For Gal80^ts^ experiments, crosses were maintained at 18°C until larvae reached late second or early third instar stages and shifted to 29°C for the duration specified in each experiment. Immediately after incubation at 29°C, wing imaginal tissues were dissected from wandering third instar larvae and imaged live.

### Live imaging

Throughout the study (except in Figs. 2A-C’, 4A-D’, S1A-A’, S5A-D’), wing imaginal discs were imaged live. Imaging of live tissues was performed as previously described (Xu et al., 2019). Briefly, freshly dissected wing imaginal discs from third instar larvae were pipetted into a ∼40 ml droplet of Schneider’s Drosophila Medium supplemented with 10% fetal bovine serum and mounted on a glass slide. To support the tissue, spherical glass beads (Cospheric, Product ID: SLGMS-2.5) of ∼50 mm in diameter were placed under the cover slip. The mounted samples were immediately imaged.

For osmotic shifts, Y-27632 treatment, and laser ablation experiments, live imaging method was adapted from Restrepo et al. (2016). Briefly, wing imaginal tissues were first dissected in Schneider’s Drosophila Medium (Sigma) supplemented with 10% Fetal Bovine Serum (Thermo Fisher Scientific). The tissues were then transferred with a pipette in 5-10μl of medium to a glass bottom microwell dish (MatTek, 35mm petri dish, 14mm microwell) with No. 1.5 coverglass. The discs were oriented so that the apical side of the disc proper faced the coverglass. A Millicell culture insert (Sigma, 12mm diameter, 8μm membrane pore size) was prepared in advance by cutting off the bottom legs with a razor blade and removing any excess membrane material around the rim of the insert. The insert was carefully placed into the 14mm microwell space, directly on top of the drop containing properly oriented tissues. To prevent the tissue from moving, the space between the insert and the microwell was sealed with ∼10μl of mineral oil. Media with indicated osmolarity and/or chemical inhibitor was then added into the chamber of the insert (200μl in all experiments). An inverted Zeiss LSM880 laser scanning confocal microscope equipped with a GaAsP spectral detector and the Airyscan module was used for all imaging (except for laser ablation experiments, see below).

### Osmotic shift experiments

Schneider’s Drosophila Medium (Sigma) supplemented with 10% Fetal Bovine Serum (Thermo Fisher Scientific) was used as isotonic medium (∼360mOsm). To make a hypertonic solution, the osmolarity of the isotonic solution was increased to ∼460mOsm using 1M NaCl. To make a hypotonic solution, the isotonic medium was diluted with deionized water to ∼216mOsm. All osmotic solutions were prepared fresh immediately before the experiments. In most experiments, tissues were incubated for 15 min in a humid chamber before mounting (see above). For practical reasons, in laser ablation experiments tissues were incubated for 8 min to allow for measurements to be taken at an average of 15 min under osmotic incubations.

To observe changes in Yki-YFP localization, flies expressing BAC recombineered Yki-YFP in the background of *yki^B5^* null allele were used. Ubi>RFP^nls^ was also expressed in the background to mark the nuclei. Tissues were dissected and mounted as described above. Each wing disc was first imaged under isotonic conditions, with 4-5 optical sections taken in the basal plane where the nuclei were clearly identifiable. The isotonic medium was then replaced with an indicated osmotic solution, and after 30 min of incubation, each tissue was reacquired in the same order and roughly the same optical plane. The steps were repeated for the third osmotic condition.

### pMRLC staining experiments

Wandering third instar larvae were dissected in an isotonic solution and the wing imaginal discs were transferred into a 150μl drop of freshly prepared osmotic medium. For experiments with Y-27632, osmotic media with 50μM of Y-27632 were prepared. The tissues were incubated for 15 min. After incubation, the discs were washed with 1X Ringer’s buffer with correspondingly adjusted osmolarity (i.e. for hypertonic experiments, a hypertonic wash was prepared the same way as described above, except 1X Ringer’s was used instead of the Schneider’s medium) and fixed for 20 min in 2% paraformaldehyde/1X Ringer’s solution also with properly adjusted osmolarity (for Y-27632 experiments, 50μM of Y-27632 was also added in the wash and the fix solutions). Tissues were then stained with the primary antibody against the *Drosophila* phospho-MRLC (anti-pSqh, see Table 1) and secondary antibodies as previously described (McCartney and Fehon, 1996).

**Table 1:** Reagents.

| Reagent type (species) or resource | Designation | Source or reference | Identifiers | Additional information |
| --- | --- | --- | --- | --- |
| genetic reagent (*D. melanogaster*) | *Ubi>Kib-GFP-FLAG* | (Tokamov et al., 2021) |  |  |
| genetic reagent (*D. melanogaster*) | *Ubi-Kib<sup>ΔWW1</sup>-GFP-FLAG* | (Tokamov et al., 2021) |  |  |
| genetic reagent (*D. melanogaster*) | *UASp-T7-RhoGEF2* | Bloomington Drosophila Stock Center | BL9387 |  |
| genetic reagent (*D. melanogaster*) | *UAS<sup>t</sup>-myr-Yki* | 10.1016/j.devcel.2018.06.017 |  |  |
| genetic reagent (*D. melanogaster*) | *UAS-Strn RNAi* | Bloomington Drosophila Stock Center | BL26736 | Validated in 10.1016/j.devcel.2018.06.017 |
| genetic reagent (*D. melanogaster*) | *UAS-Par-1 RNAi* | Bloomington Drosophila Stock Center | BL32410 |  |
| genetic reagent (*D. melanogaster*) | *UAS-jub RNAi* | Bloomington Drosophila Stock Center | BL30806 | Validated in 10.1016/j.cub.2010.02.035 |
| genetic reagent (*D. melanogaster*) | *Ecad-3XmKate2* | (Pinheiro et al., 2017) |  |  |
| genetic reagent<br>(*D. melanogaster*) | *Par-1:GFP*<br>(protein trap) | Bloomington<br>Drosophila Stock<br>Center | BL64452 |  |
| genetic reagent<br>(*D. melanogaster*) | *Jub:GFP* | Bloomington<br>Drosophila Stock<br>Center | BL56806 |  |
| genetic reagent<br>(*D. melanogaster*) | *w<sup>1118</sup>; yki<sup>B5</sup>*<br>{*yki-YFP*}<br>*VK37* | (Xu et al., 2018) |  |  |
| genetic reagent<br>(*D. melanogaster*) | *aPKC<sup>as4</sup>* | (Hannaford et al.,<br>2019b) |  |  |
| Antibody | anti-pSqh<br>(Guinea pig,<br>polyclonal) | (Zhang and<br>Ward, 2011) |  | Tissue<br>staining<br>(1:300) |
| Antibody | anti-DEcad<br>(Rat,<br>polyclonal) | Developmental<br>Studies<br>Hybridoma Bank | Cat#AB_5<br>28120;<br>RRID:AB<br>_528120 | Tissue<br>staining<br>(1:1000) |
| Antibody | anti-PKC,<br>rabbit<br>polyclonal | Santa Cruz<br>Biotechnology | Lot#:<br>G2304 | tissue<br>staining<br>(1:1000) |
| Antibody | anti-GFP,<br>guinea pig,<br>polyclonal | (Yee et al., 2019) |  | IP (1:1250) |
| Antibody | anti-GFP,<br>rabbit,<br>polyclonal | Michael Glotzer,<br>(University of<br>Chicago) |  | IB (1:5000) |
| Chemical | Y-27632 | Fisher Scientific | Cat. # 12541 | Rho kinase<br>inhibitor |
| Chemical | 1-NA-PP1 | Cayman | Cat. #<br>CAYM-10954-1 | ATP analog |

### Laser ablation of bi-cellular junctions

Laser cuts were conducted using a pulsed Micropoint nitrogen laser (Andor Technology) tuned to 365 nm and mounted on an Andor Revolution XD spinning disk confocal microscope. Individual junctions were ablated by delivering 3 pulses at a single point with a duration of 67 ms/pulse. Each tissue was imaged 2x before and 20x immediately after ablation, with a time interval of 4 s. Images were acquired using a 100X oil-immersion lens (Olympus, NA 1.40), an iXon Ultra 897 camera (Andor), and Metamorph (Molecular Devices) as the image acquisition software. Sixteen-bit Z-stacks were collected at each time point consisting of 7 slices with 0.5 μm interval.

To measure initial recoil velocity, the positions of the two tricellular vertices connected by the ablated junction were manually tracked using the SIESTA image analysis platform (Fernandez-Gonzalez and Zallen, 2011), and the initial retraction velocities were calculated using custom MATLAB scripts.

### Image analysis, quantification of fluorescence intensities and apical areas

All images were processed in ImageJ. For apical Kib-GFP, Kib^ΔWW1^-GFP, Jub-GFP, and Ecad-mKate2 mean fluorescence intensity measurements were taken from maximum intensity projections (∼0.75 μm/section, four to six sections from apical surface). For basal Kib-GFP mean fluorescence intensity measurements, single basal optical sections were used (∼7.5-10μm below the apical surface). To quantify changes in pMRLC intensity, mean fluorescence measurements were obtained from single most apical sections. Plots and statistical analyses of mean fluorescence intensities were generated using GraphPad Prism software.

To measure changes in apical cell areas under osmotic shifts, cells from Ecad-GFP labeled epithelia were segmented using the Cellpose v1 pre-trained "cytoplasm" model (Stringer et al., 2021) and tracked via a previously-described tracking algorithm (Williams et al., 2022). Errors in segmentation and tracking were corrected manually. Only cells that remained in the field of view for the entire timelapse duration were used for quantification. The mean and standard error in cell areas were measured per epithelium. Fold-change in area relative to the start of osmotic shift was plotted using the Matplotlib library in Python.

To quantify nuclear/cytoplasmic Yki-YFP, single optical sections were used. Ubi>RFP channel was used to segment the nuclei using Cellpose and a standard Scikit watershed algorithm (van der Walt et al., 2014) to generate a nuclear mask. The nuclear mask was then applied to Yki-YFP images to extract nuclear Yki-YFP intensities, and the signal outside the nuclear mask was treated as cytoplasmic. The ratios of nuclear to cytoplasmic intensity was then calculated and plotted.

### Detection of Kibra ubiquitination in S2 cells

Kib ubiquitination assay was performed as described previously (Tokamov et al., 2021). Briefly, 3.5 x 10^6^ S2 cells (S2-DGRC) were transfected with a total of 500 ng of indicated DNA using dimethyldioctadecylammonium bromide (Sigma; Han, 1996) at 250 mg/ml in six-well plates. pMT-Kib-GFP was co-transfected with pMT-HA-Ub (Zhang et al., 2006) where indicated to provide labeled ubiquitin. To induce expression of the pMT constructs, 700 mM CuSO4 was added to the wells 24 hr prior to cell lysis (2 days after transfection). To inhibit proteasomal degradation, 50 mM MG132 (Cayman Chemical) and 50 mM calpain inhibitor I (Sigma Aldrich) was added 4 hr prior to cell lysis. Cells were lysed in RIPA buffer (150 mM NaCl, 1% NP-40, 0.5% Na deoxycholate, 0.1% SDS, and 25 mM Tris [50 mM, pH 7.4]), supplemented with 5 mM N-ethylmaleimide and Complete protease inhibitor cocktail (Roche, one tablet/10 ml of buffer). HA-tagged ubiquitin was purified using Pierce anti-HA magnetic beads (clone 2–2.2.14). Lysates and IP samples were run on 8% polyacrylamide gel and ubiquitinated Kib in the IP samples was detected by Western Blot using anti-GFP antibody (rabbit, see Table 1).

### Detection of Kibra phosphorylation in S2 cells

pMT-Kib-GFP and pAHW-Par-1 or pAHW-Par-1^KD^ (gift from Bingwei Lu, Stanford University) were transfected and expression was induced as described above. Immunoprecipitation (IP) was performed 3 days after transfection. To induce expression of pMT-Kib-GFP, 700 mM CuSO4 was added to the wells 24 hr prior to cell lysis (2 days after transfection).

Cells were harvested and lysed on ice in buffer containing 25 mM Hepes, 150 mM NaCl, 1 mM ethylenediaminetetraacetic acid, 0.5 mM ethylene glycol-bis(b-aminoethyl ether)-N,N,N0,N0-tetraacetic acid, 0.9 M glycerol, 0.1% Triton X-100, 0.5 mM Dithiothreitol, and Complete protease inhibitor (Roche) and PhosSTOP (Sigma Aldrich) phosphatase inhibitor cocktails at one tablet/10 ml concentration each. Cell lysates were then incubated with anti-GFP antibody (guinea pig, see Table 1) for 30min. Antibody-bound Kib-GFP was pulled down using Pierce Protein A magnetic beads (Thermo Scientific) for 1.5h. A control immunoprecipiated sample was treated with λ-phosphatase. Samples were run on 8% polyacrylamide gel, with 118:1 acrylamide/bisacrylamide (Scheid et al., 1999), to better resolve phosphorylated Kib species. Kib was detected by Western Blot using anti-GFP antibody (rabbit, see Table 1).

