## Supplemental figures and legends for "Cortical tension regulates Hippo signaling via Par-1-mediated Kibra degradation"

### Figure S1, related to Fig. 1

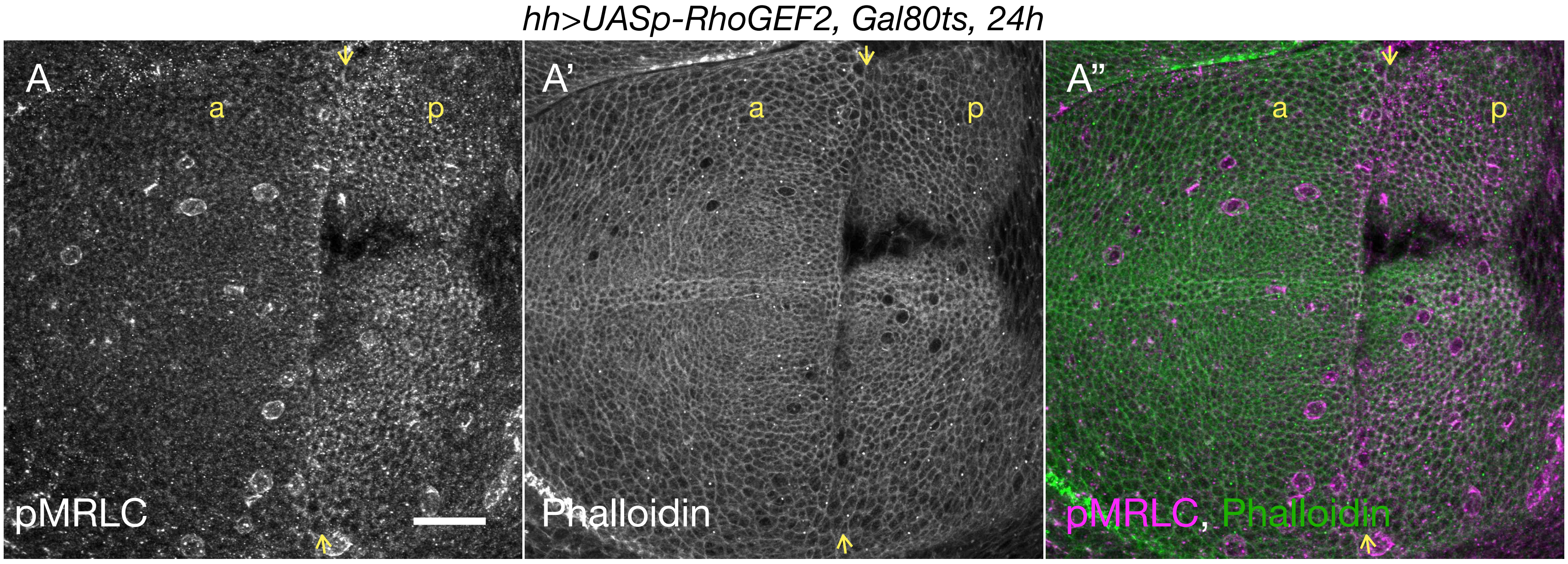


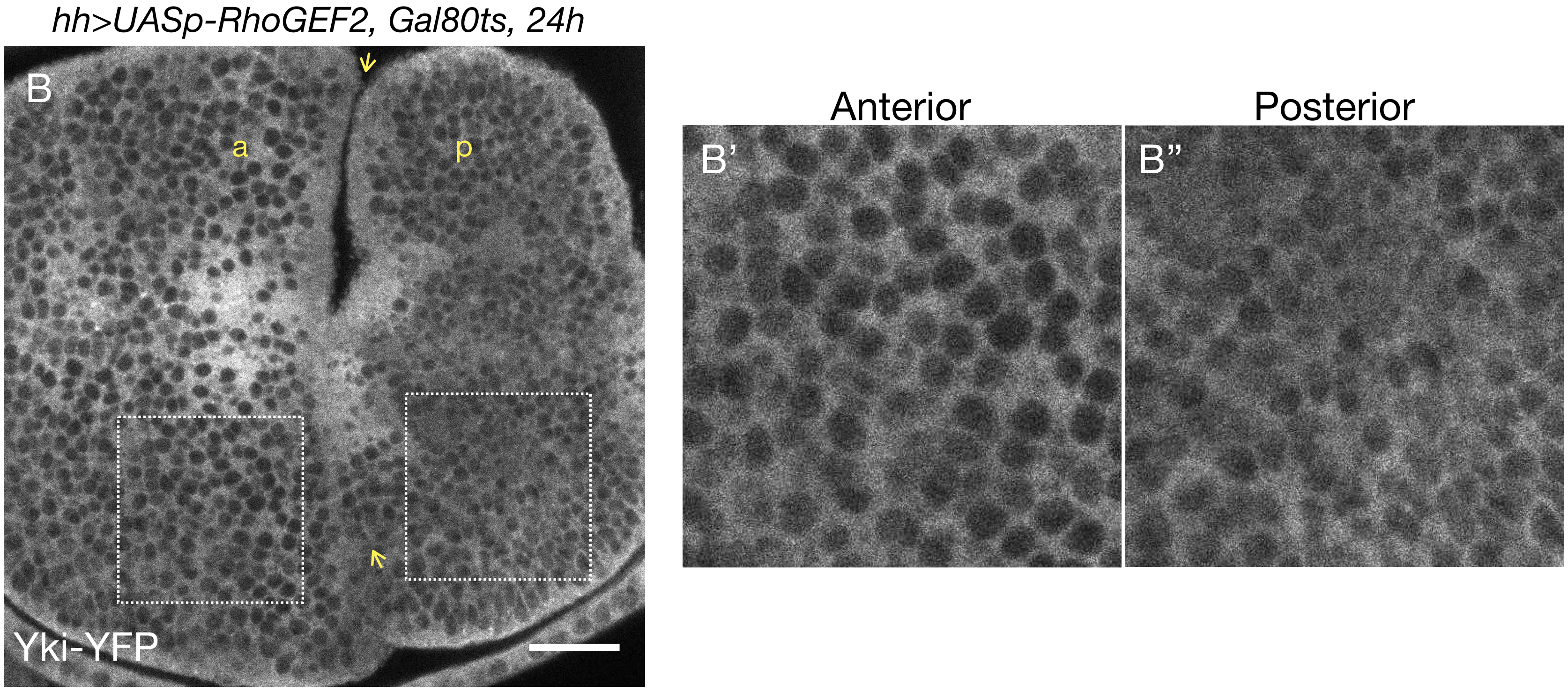

**Figure S1: Transient ectopic RhoGEF2 expression increases myosin activation and nuclear Yki accumulation.**

A-A'') A representative image of a wing imaginal disc ectopically expressing RhoGEF2 in the posterior compartment for 24h and stained for pMRLC (A) and F-actin (phalloidin, A'). Yellow arrows indicate the anterior-posterior (a-p) boundary. Note the increase in pMRLC in the posterior (p) compartment.

B-B'') Yki-YFP becomes more nuclear in the posterior compartment, where RhoGEF2 was ectopically expressed for 24h.

Scale bars = 20μm.

Figure S2, related to Fig. 2

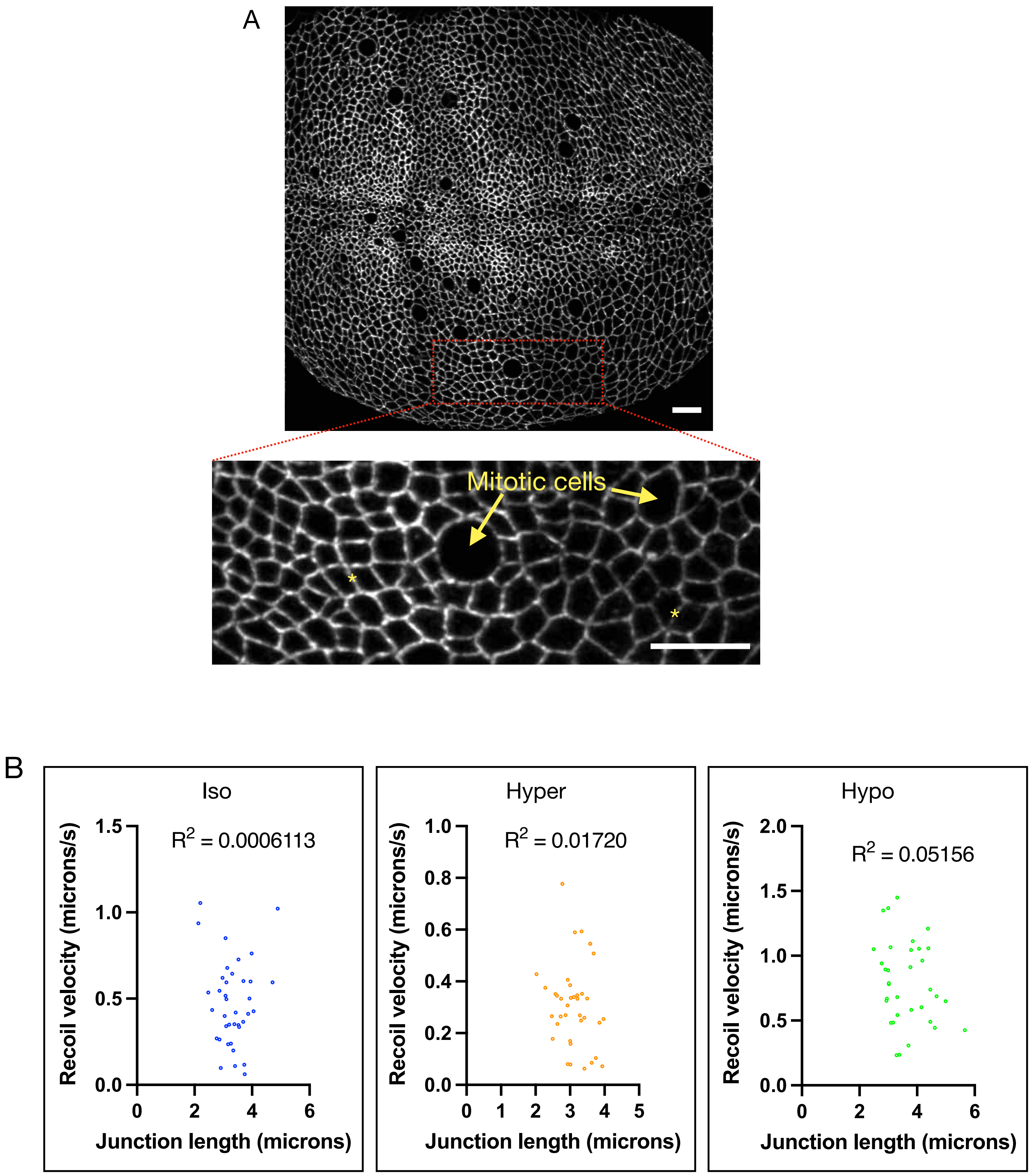

**Figure S2: Laser ablations of bicellular junctions under osmotic shifts.**
A) An example of a wing imaginal tissue expressing Ecad-mKate2 as a junctional marker used for laser cutting experiments. The enlarged region indicates an approximate area (anterior ventral region) where junctional cuts were generated. No more than two junctions (yellow asterisks) were cut per tissue. The ablated junctions were multiple cells apart and never in direct contact with mitotic cells. Scale bars = 10μm.
B) There was no relationship between initial junction length and recoil velocity. The $R^2$ values represent Pearson correlation coefficients.

### Figure S3, related to Fig. 2

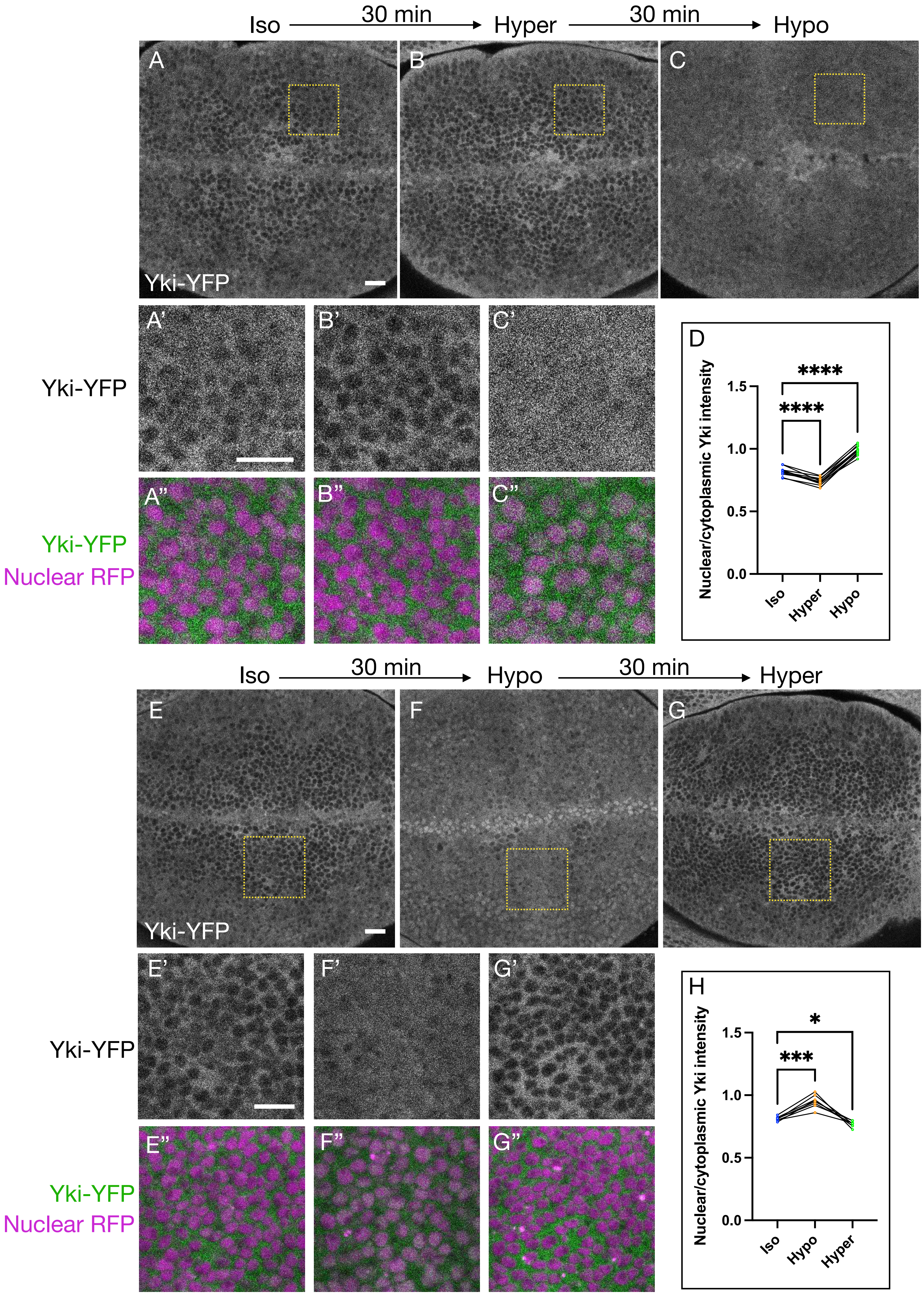

**Figure S3: Effect of osmotic shifts on nuclear Yki localization.**
A-C'') Compared to isotonic conditions (A-A''), Yki becomes less nuclear in hypertonic medium (B-B'') and more nuclear in hypotonic medium (C-C''). Note that the same tissue is displayed in A-C.
D) Quantification of nuclear/cytoplasmic mean Yki-YFP fluorescence under osmotic conditions represented in A-C''.
E-G'') Similar experiment as shown in A-C'', except tissues were first shifted from isotonic (E-E'') to hypotonic (F-F'') medium to make the effect of hypertonic conditions more evident. Shifting from hypotonic to hypertonic conditions resulted in a dramatic decrease of nuclear Yki (G-G''). Note that the same tissue is displayed in E-G. Scale bars = 10μm.
H) Quantification of nuclear/cytoplasmic mean Yki-YFP fluorescence under osmotic conditions represented in E-G''. Statistical significance in D & H was calculated using Repeated-measures One-way ANOVA test followed by Tukey's HSD test.

### Figure S4, related to Fig. 3

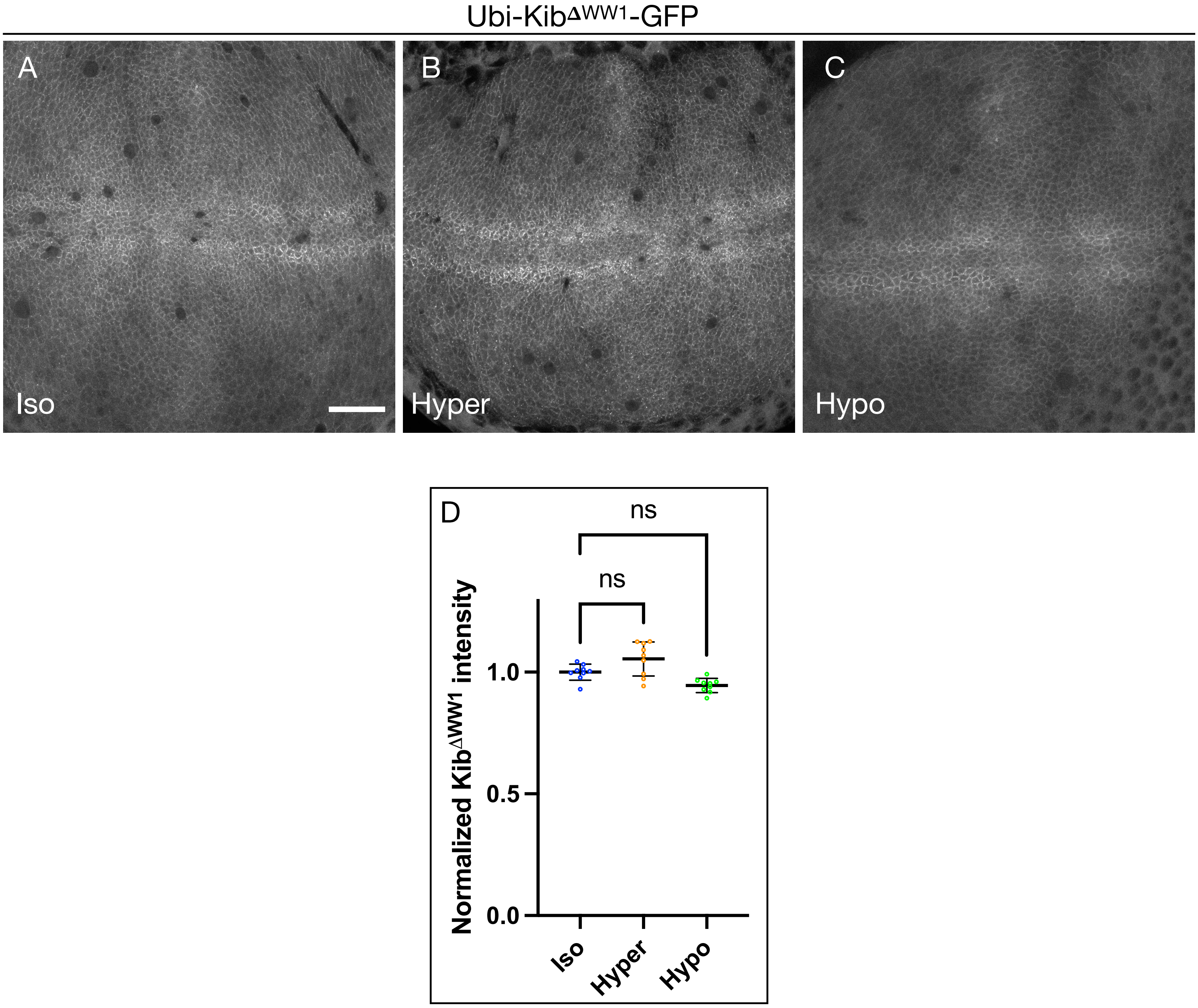

**Figure S4: Osmotic shifts have no significant effect on Kib$^{\Delta WW1}$ abundance.**
A-C) Representative wing imaginal tissues expressing Ubi-Kib$^{\Delta WW1}$-GFP incubated under isotonic (A), hypertonic (B), and hypotonic (C) conditions. Scale bar = 20µm.
D) Plot of normalized Ubi-Kib $^{\Delta WW1}$-GFP mean intensities under osmotic conditions shown in A-C.

### Figure S5, related to Fig. 4

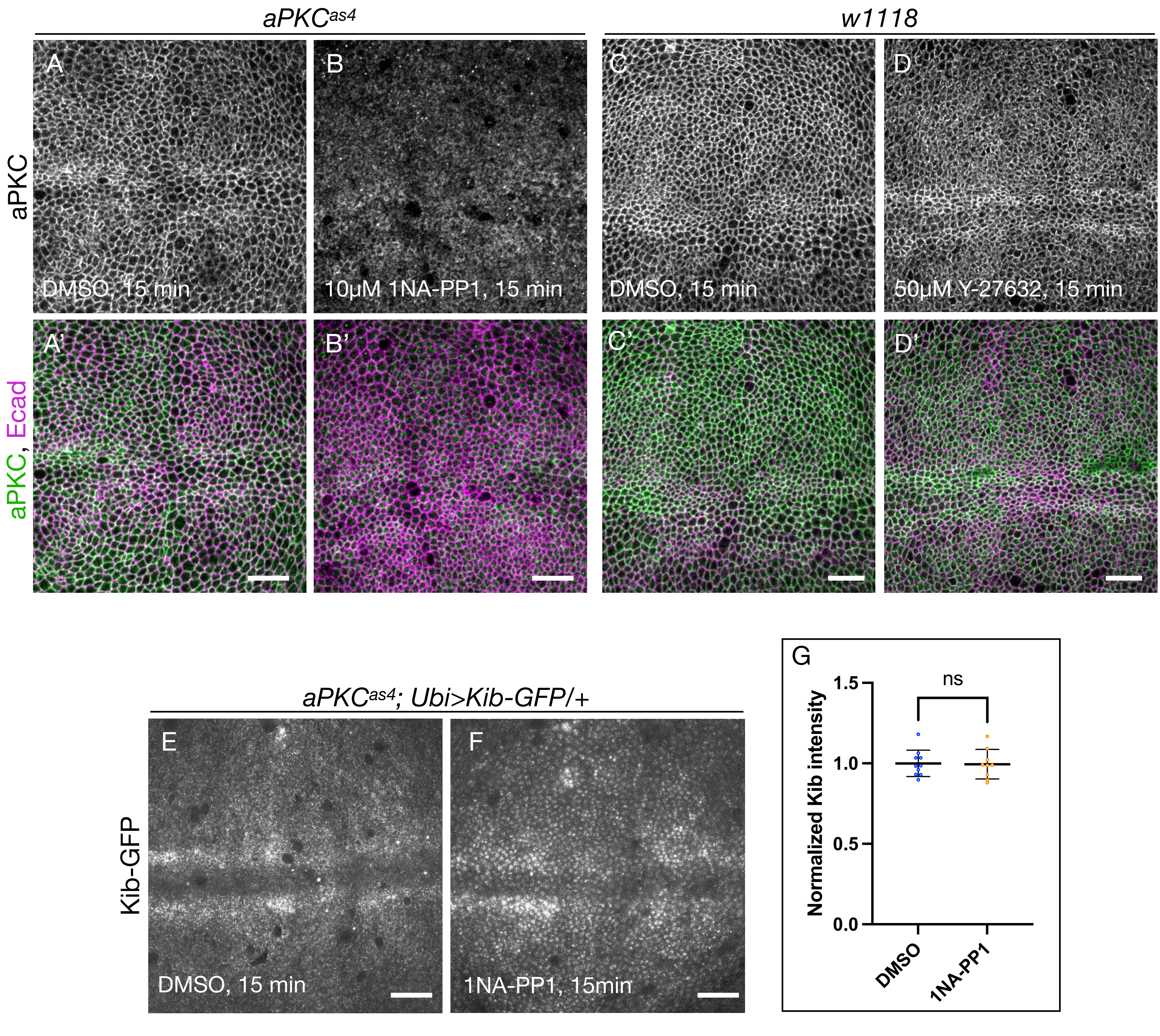

**Figure S5: aPKC inhibition does not affect Kib abundance.**
A-B') aPKC and Ecad immunostaining shows that while $aPKC^{as4}$ is normally cortically localized in DMSO-treated wing discs (A & A'), this localization was significantly inhibited upon treatment with 1NA-PP1 (B & B'). Scale bars = 10µm.
C-D') In wild type tissues aPKC localizes cortically under DMSO treatment (C & C'). Treatment with Y-27632 does not significantly affect cortical aPKC localization (D & D'). Scale bars = 10µm.
E-G) In the homozygous $aPKC^{as4}$ background, Kib abundance was similar in DMSO-treated (E) and 1NA-PP1-treated tissues (F). Scale bars = 10µm. Statistical significance was calculated using Mann-Whitney test.

### Figure S6, related to Fig. 5

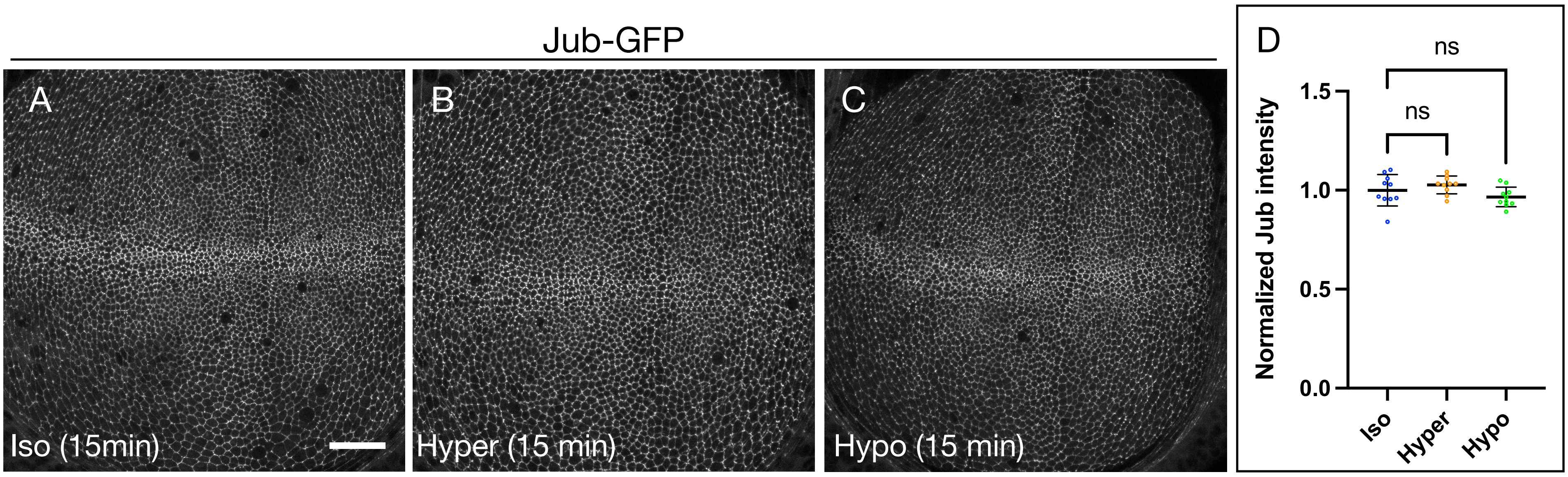


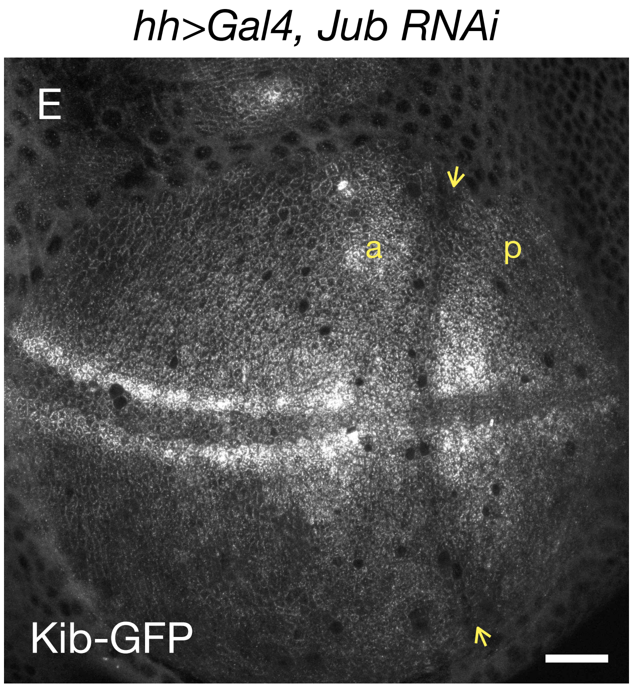

**Figure S6: Tension-mediated Kib degradation is independent of the Jub-Wts mechanism and is regulated via Par-1.**

A-C) Representative wing imaginal tissues expressing Jub-GFP incubated under isotonic (A), hypertonic (B), and hypotonic (C) conditions.

D) Plot of normalized Jub-GFP mean intensities under osmotic conditions shown in A-C. Statistical significance was calculated using One-way ANOVA followed by Tukey's HSD test.

E) Depletion of Jub in the posterior compartment of the wing disc does not affect Kib levels. Yellow arrows indicate the anterior-posterior (a-p) boundary. Scale bars = 20µm.

### Figure S7, related to Fig. 5

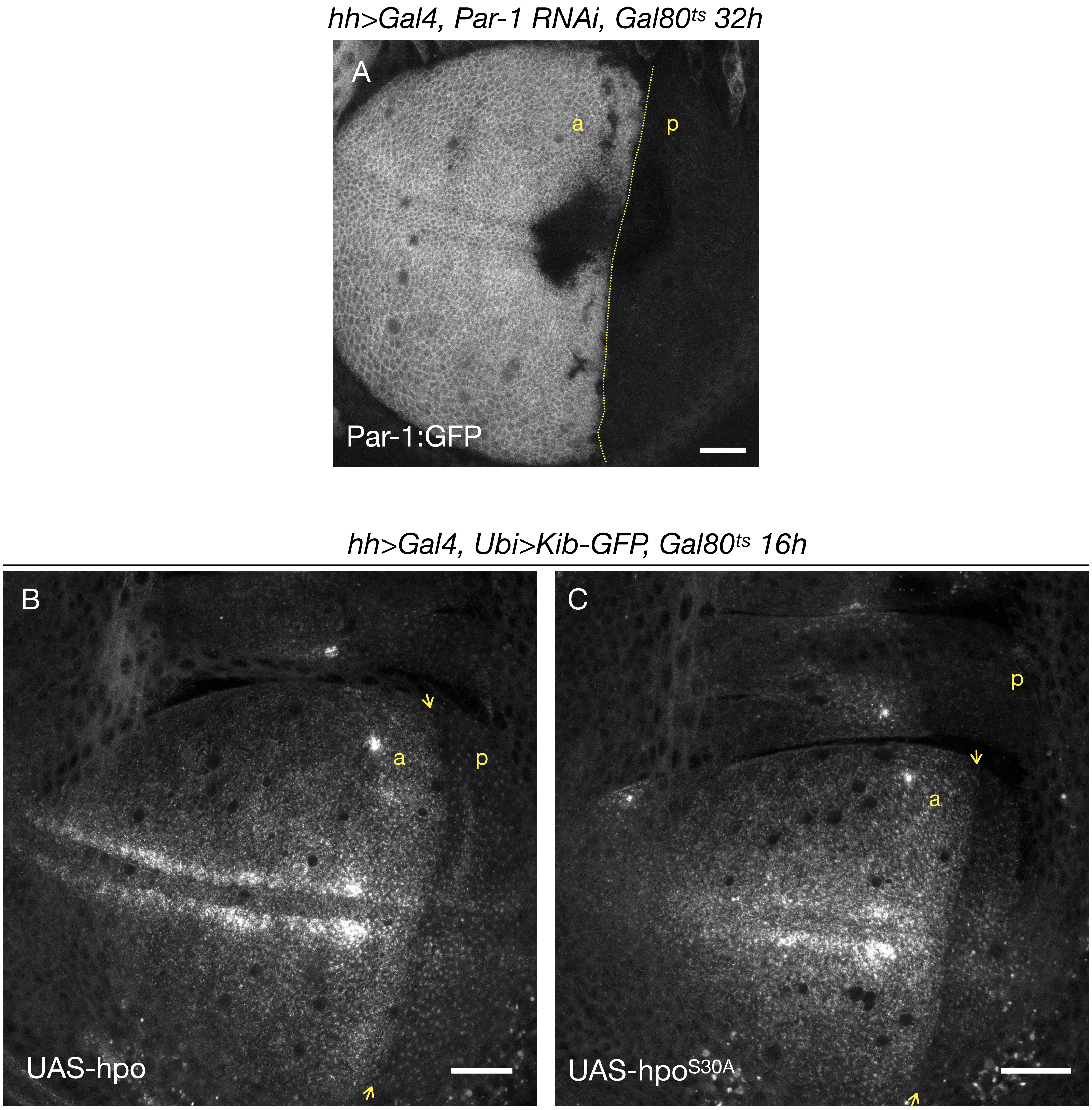

**Figure S7: Par-1 regulates Kib abundance independently of Hpo phosphorylation on Ser30.**

A) Validation of Par-1 depletion. A short hairpin RNA targeting Par-1 was expressed for 32h in the posterior compartment (p) of the wing disc expressing a GFP-trap of Par-1. Yellow line indicates the anterior-posterior (a-p) boundary.

B-C) Transient ectopic expression of either wild-type Hpo (B) and $Hpo^{S30A}$ (C) in the posterior compartment (p) of the wing imaginal disc leads to similar decrease in Kib abundance. Yellow arrows indicate the anterior-posterior (a-p) boundary. Scale bars = 20μm.
